# Carbs versus fat: does it really matter for maintaining lost weight?

**DOI:** 10.1101/476655

**Authors:** Kevin D. Hall, Juen Guo

## Abstract

- The latest battle in the perpetual diet wars claimed that low carbohydrate diets offer a metabolic advantage to burn more calories and thereby help patients maintain lost weight.
- However, analyzing the data according to the original pre-registered statistical plan resulted in no statistically significant effects of diet composition on energy expenditure.
- The large reported diet effect on expenditure calculated using the revised analysis plan depended on data from subjects with excessive amounts of unaccounted energy. Adjusting the data to be commensurate with energy conservation resulted in a diet effect that was less than half the value reported in the BMJ paper.
- The measured daily average CO_2_ production rates were not significantly different between the diets and the reported expenditure differences were due to inaccurate calculations based on false assumptions about diet adherence.

---

Proponents of low carbohydrate diets claim that they result in greater calorie expenditure thereby providing patients with a “high calorie way to stay thin forever” ^1^. While most studies have found no clinically meaningful effect of low carbohydrate diets on total energy expenditure (TEE) ^2^, it’s possible that prior studies simply failed to create the appropriate conditions to reveal the effect. For example, perhaps low carbohydrate diets help prevent the usual reduction in TEE that occurs after losing weight – an effect that may help people avoid weight regain. In support of this possibility, Ebbeling et al. concluded that low carbohydrate diets substantially increase TEE compared to high carbohydrate diets during maintenance of lost weight ^3^. Unfortunately, the data may not support this conclusion.

## Reported data analysis was not conducted according to the original plan

Registering a clinical trial’s primary outcome and statistical analysis plan *before data is collected* helps reduce bias and improves scientific reproducibility ^4^. The original pre-registered protocol and analysis plan of Ebbeling et al. addressed whether the *reduction* in TEE during maintenance of lost weight depended on the dietary carbohydrate to fat ratio *when compared to the pre-weight loss baseline* – a design similar to a pilot study by the same authors ^5^ whose data were used by to design and statistically power the new study. The pre-registered plan was in place for most of the study’s history, including 7 of 8 protocol versions between 2014-2016.

However, the analysis plan was modified in 2017 after all subjects had completed the trial and after primary data for the first two of three cohorts were returned to the unblinded principal investigators in Boston from the blinded doubly labeled water (DLW) laboratory in Houston. Nevertheless, according to the principal investigator, the Boston statistician who performed the data analyses was unblinded to the diet assignments only after all primary data were returned from the Houston lab and after the revised analysis plan was registered. The change in analysis plan was not acknowledged in the original manuscript submission to The BMJ and was not reported in a previous publication of the trial design ^6^.

The revised primary outcome compared TEE during weight loss maintenance to TEE measured in the immediate post-weight loss period rather than the originally planned pre-weight loss baseline. No reasons for the change were provided in the final protocol or statistical analysis plan. The BMJ publication stated the original plan was an “error” and their Data Supplement listed three reasons for the change. First, post-weight loss TEE was closer to the time of diet randomization. Second, *pre-weight loss* TEE was “strongly confounded by weight loss”. How this might happen is difficult to imagine. Finally, the original plan was claimed to be under-powered despite the study being designed and its power calculations being informed by pre-weight loss TEE data of the pilot study that did not measure TEE in the period immediately post-weight loss ^5^. Interestingly, Ebbeling et al. justified the claim that the original plan was underpowered using a post hoc analysis showing that the original plan did not result in a significant diet effect.

The original plan was preferable for several reasons. First, it was the only plan registered before data were collected. Second, it addressed the question of whether the *reduction* in TEE that accompanies maintenance of lost weight depends on the dietary carbohydrate to fat ratio. Third, the pre-weight loss baseline DLW measurements were obtained in the routine situation when people were maintaining their habitual weight. In contrast, the revised plan relied on post-weight loss measurements that were obtained during the period when diet calories were increasing at a rate determined by each individual subject’s recent rate of weight loss. The DLW method has never been validated in such a refeeding condition which introduces uncertainty into the TEE calculations because the respiratory quotient (RQ) was certainly not equal to the food quotient as assumed by Ebbeling et al. ^7^. Furthermore, TEE measurements in the immediately post-weight loss period were potentially confounded by transient adaptive thermogenesis that typically becomes less severe after an extended weight stabilization period ^8 9^. Therefore, post-weight loss DLW measurements should ideally have been conducted after subjects had stabilized at the lower body weight for several weeks while consuming a constant diet.

## The original pre-registered analysis plan did not result in significant diet differences

Despite the BMJ Editors’ request to report the results of their original analysis plan, Ebbeling et al. argued against this because they were “concerned that the additional analysis would provide no meaningful biological insights – that is, no useful information about the nature of the relationship between dietary composition and energy expenditure.” Here, we report the results of the originally planned analyses while deviating as little as possible from the methods published by Ebbeling et al. ^3^.

We downloaded the individual subject data and SAS statistical analysis code from the Open Science Framework website (https://osf.io/rvbuy/) and modified the code to reanalyze the data according to the original plan. Because Ebbeling et al. claimed that the per protocol group “provide a more accurate estimate of the true diet effects”, we focus our attention on this group and report the intention to treat analysis in the Appendix.

When using the original analysis plan, we found no significant diet differences. Pairwise TEE comparisons with respect to the pre-weight loss baseline were not significant between diets (p>0.35) (Figure 1A). The low, moderate, and high carbohydrate groups decreased TEE by (mean ± SE) 262 ±72 kcal/d, 254±75 kcal/d, and 356±80 kcal/d, respectively, compared with the pre-weight loss baseline period (p=0.59 for the test of equivalence between the diets). The linear trend estimate was 24±27 kcal/d per 10% decrease in carbohydrate (p=0.38). The mean absolute weight losses at 10 and 20 weeks compared to the pre-weight loss baseline were well-matched and within 250 g between all diet groups (p>0.9), so any diet effects could not have been obscured by group differences in mean weight loss. Similar results were obtained when using weight-normalized TEE.

**Figure 1.**
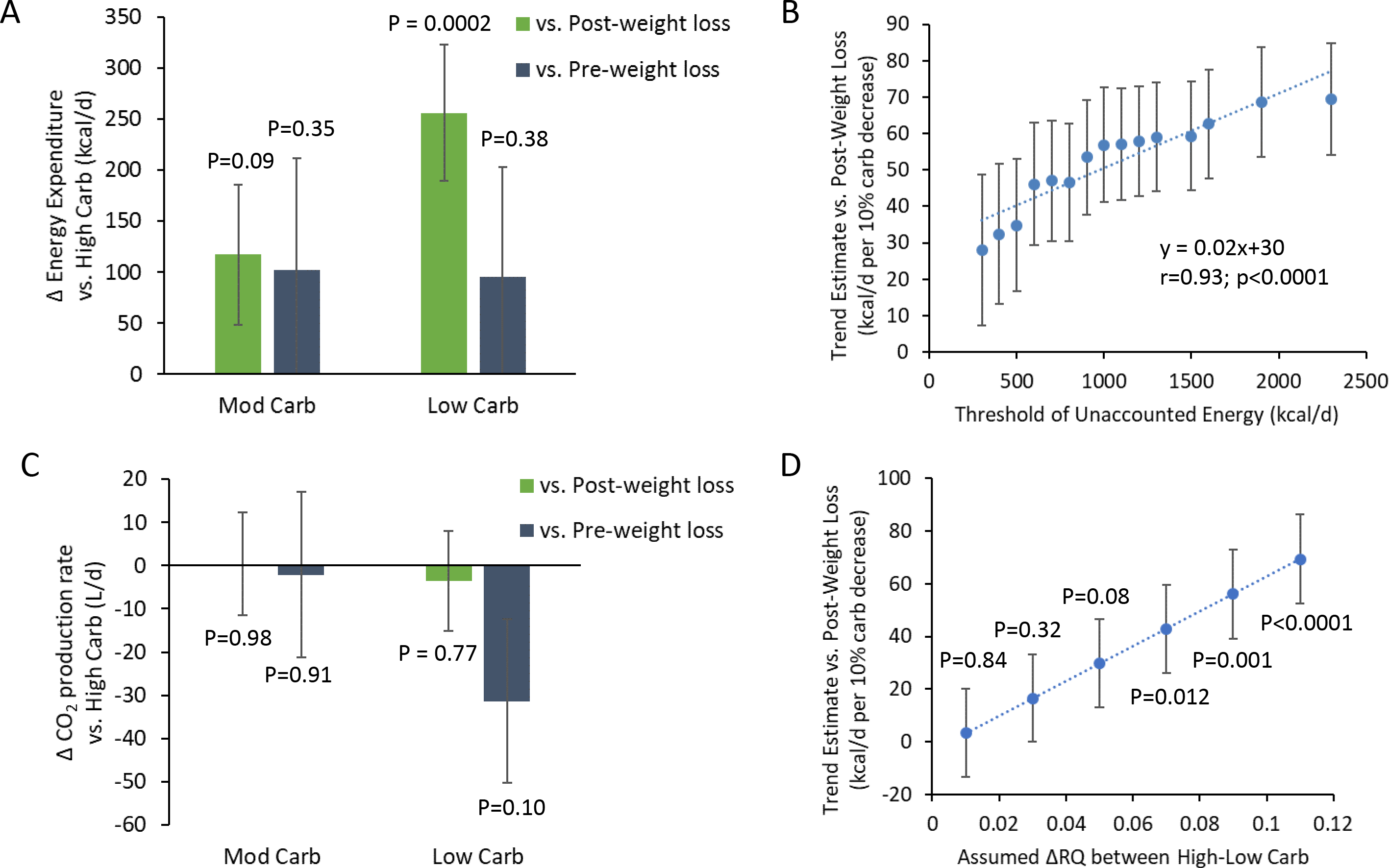
A) Differences in total energy expenditure (TEE) in the per-protocol group consuming low and moderate carbohydrate diets compared to subjects consuming a high-carbohydrate diet. The gray bars indicate the lack of significant effect of diet on average TEE during weight loss maintenance as compared to the pre-weight loss baseline period according to the original analysis plan of Ebbeling et al. The green bars illustrate how the revised analysis plan resulted in a significant effect of the low carbohydrate diet on average TEE during weight loss maintenance as compared to the immediate post-weight loss period. B) Per-protocol trend estimate for the TEE diet effect during weight loss maintenance (using the revised analysis plan) as a function of the threshold used to filter out subjects with excessive amounts of unaccounted energy. The rightmost data point includes all 120 per-protocol subjects with as much as 2300 kcal/d of unaccounted energy and corresponds to the diet effect size reported by Ebbeling et al. according to their revised analysis plan. The leftmost data point indicates a reduced effect size and includes 57 subjects with as much as 300 kcal/d of unaccounted energy. C) Differences in daily average CO_2_ production in the per-protocol group consuming low and moderate carbohydrate diets compared to subjects consuming a high-carbohydrate diet. No significant effects of diet were observed regardless of whether the measurements during weight loss maintenance were compared to the pre-weight loss baseline period (gray bars) or compared to the immediate post-weight loss period (green bars). D) Per-protocol trend estimate for the TEE diet effect during weight loss maintenance (using the revised analysis plan) as a function of the assumed differences in daily RQ between the high and low carbohydrate diets. Error bars are ±SE.

## Reported diet differences were inflated by subjects with implausible unaccounted energy

Although Ebbeling et al. provided the subjects with all their food to maintain a stable lower body weight, the reported energy intake was 422 ± 47 kcal/d (p<0.0001) less than TEE. Such large apparent energy deficits during a period of relative weight stability indicate that the subjects were likely consuming a substantial amount of unaccounted food and beverages despite the controlled-feeding design.

The law of energy conservation requires that accurate measurements of energy intake, TEE, and body weight change be quantitatively commensurate. Unfortunately, the data of Ebbeling et al. revealed extraordinary amounts of unaccounted energy in many subjects. Because body composition measurments have yet to be reported, we assumed that each kg of weight change corresponded to an energy content of 7700 kcal. We calculated that 483 ± 39 kcal/d (p<0.0001) were unaccounted during the weight loss maintenance period (see the Appendix). Such large amounts of missing calories, in violation of the law of energy conservation, indicates that the measurements of intake and TEE were inaccurate and imprecise for many subjects.

Importantly, the large reported TEE diet effects according to the revised analysis plan depended on including subjects with excessive unaccounted energy. Figure 1B illustrates the significant attenuation of the diet effect when increasingly stringent thresholds were employed to remove subjects with excessive amounts of unaccounted energy (r=0.93; p<0.0001). The intercept of the best fit line was 30±3 kcal/d per 10% reduction in dietary carbohydrates and corresponds to the estimated diet effect on TEE when all energy is accounted. In other words, the TEE diet effect was less than half the value reported by Ebbeling et al. after adjusting the data to be commensurate with the law of energy conservation.

## Diet differences in subjects with high insulin secretion were uncorroborated by other measures

The significant ^~^500 kcal/d effect modification of TEE by baseline insulin secretion observed by Ebbeling et al. when using the post-weight loss TEE measurement as the anchor point was no longer significant when using the pre-weight loss TEE as the anchor point (p=0.36 for the test of equivalence between the diets). Nevertheless, for subjects in the highest insulin secretion tertile TEE was 383±196 kcal/d greater for the low versus high carbohydrate diets (p=0.053). While not as large as the reported effect size using the revised analysis plan, such TEE differences in the high insulin secretion group could be physiologically important.

Was this potential TEE effect corroborated by corresponding differences in measured components of energy expenditure? Unfortunately, it was not. Differences in resting energy expenditure (−32±49 kcal/d; p=0.52), total physical activity (45754±47821 counts/d; p=0.34), moderate to vigorous physical activity (−5±6 min/d; p=0.4), sedentary time (−9±30 min/d; p=0.77), skeletal muscle work efficiency at 10W (1±0.9 %; p=0.27), 25W (1.2±1.1 %; p=0.28) and 50W (0.5±0.8 %; p=0.48) were all not significantly different between the low and high carbohydrate diets when compared to the pre-weight loss baseline. Nevertheless, we cannot rule out possible differences in the thermic effect of food, sleeping energy expenditure, or another unmeasured factor contributing to TEE. Alternatively, the apparent TEE diet differences may have been due to inaccurate DLW calculations ^7^.

## Daily average CO_2_ production was not different between the diets

Using the TEE data along and the DLW equations reported by Ebbeling et al., we back-calculated the DLW measurements of daily average CO_2_ production. Figure 1C shows that there were no significant diet differences in CO_2_ production regardless of whether the pre-weight loss or post-weight loss point was used as the basis for comparison. Therefore, the reported diet differences entirely relied on the assumptions about daily RQ ^7^.

Given the large discrepancies between reported energy intake and TEE described above, along with the fact that many study meals and snacks were unsupervised and therefore their consumption was unverified, the daily RQ values were unlikely to equal the food quotients of the study diets. Nevertheless, Ebbeling et al. assumed that the differences in daily RQ between the high and low carbohydrate diets was at the theoretical maximum value of 0.11 assuming energy balance and strict diet adherence. In contrast, daily RQ differences were <0.1 when directly measured using respiratory chambers in subjects admitted to metabolic wards to ensure strict adherence to isocaloric diets varying in carbohydrates by a greater degree than those of Ebbeling et al. ^7^. Therefore, the RQ assumptions of Ebbeling et al. likely overestimated TEE differences between the low and high carbohydrate diets.

We investigated the effect of varying the assumed daily RQ differences between the high and low carb diets. Figure 1D illustrates that the TEE effect calculated using the revised post-weight loss anchor point was no longer statistically significant if the high and low carb diets resulted in daily RQ values of 0.87 and 0.82 rather than the assumed values of 0.9 and 0.79, respectively. Such differences from the assumed RQ and food quotient values are within the uncertainty range observed in an inpatient metabolic ward study with subjects strictly adhering to isocaloric diets varying widely in their carbohydrate to fat ratio ^7^.

## Conclusion

The significant effects of low carbohydrate diets on TEE reported by Ebbeling et al. failed to materialize when data were analyzed according to the original pre-specified plan. Furthermore, the significant results obtained by the revised analysis plan required unrealistic DLW calculations and required inclusion of subjects with implausibly large amounts of unaccounted energy. Therefore, in accord with many prior studies on this topic ^2^, the data of Ebbeling et al. do not support the concept that low carbohydrate diets substantially affect energy expenditure.

## Competing Interests

KDH has participated in a series of debates with Dr. David S. Ludwig, the senior author of the main study in question, regarding the merits and demerits of the carbohydrate-insulin model of obesity as well as the physiological response of the human body to isocaloric diets varying in the ratio of carbohydrates to fat.

## APPENDIX

### Intention to treat analysis

In the intention to treat analysis according to the original plan, no significant differences in TEE were found between diet groups compared with the pre-weight loss baseline period; with the low, moderate, and high carbohydrate groups decreasing TEE by 240 ±64 kcal/d, 322±66 kcal/d, and 356±67 kcal/d, respectively (p=0.43 for the test of equivalence between the diets). Pairwise comparisons of TEE diet differences with respect to the pre-weight loss baseline were not significant between diets (p>0.35) (Figure S1A). The linear trend estimate was 29±23 kcal/d per 10% decrease in carbohydrate (p=0.21). Similar results were obtained using weight-normalized TEE data.

The measured energy intake was 460 ± 46 kcal/d (p<0.0001) less than TEE. When calculated according to the revised analysis plan of Ebbeling et al., the large TEE diet effect depended on including subjects with excessive unaccounted energy. Figure S1B illustrates the significant effect size attenuation when increasingly stringent thresholds were employed to remove subjects with excessive unaccounted energy (r=0.91; p<0.0001). The intercept of the best fit line was 30±2 kcal/d per 10% reduction in dietary carbohydrates and corresponds to the estimated diet effect size when all energy is accounted. In other words, the TEE diet effect was about half the value reported by Ebbeling et al. after adjusting the data to be commensurate with the law of energy conservation.

Figure S1C shows that there were no significant diet differences in CO_2_ production regardless of whether the pre-weight loss or post-weight loss point was used as the basis for comparison. Figure S1D illustrates that the TEE effect calculated using the revised post-weight loss anchor point was no longer statistically significant if the high and low carb diets resulted in daily RQ values of 0.88 and 0.81 rather than the assumed values of 0.9 and 0.79, respectively.

### Unaccounted Energy

The law of energy conservation applied to human body weight (BW) dynamics requires that the following equality hold:

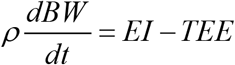

where the left side of the equation is the rate of change in body energy stores with ρ being the energy density of the weight change. On the right side of the equation, EI is the metabolizable energy intake and TEE is the total energy expenditure. EI was controlled and periodically adjusted to ensure that BW was relatively stable (i.e., the left side of the equation was approximately zero).

Unaccounted energy, UE, was defined as:

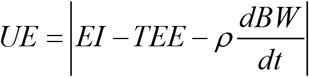

which is ideally zero. Of course, imperfect measurements will lead to nonzero values for unaccounted energy, but the values should be small and representative of the imprecision of the instruments used to make the measurements. We calculated UE from 0-10 weeks and from 10-20 weeks using the mean values of EI and TEE over each interval along with the estimated value of ρ = 7700 kcal/kg assumed by Ebbeling et al. for their EI adjustments ^10^. Ideally, the body composition measurements would have provided a more accurate assessment of changes in body energy stores, but these data have not yet been made available by Ebbeling et al. ^10^.

## Supplementary Figure Legend

**Figure S1.**
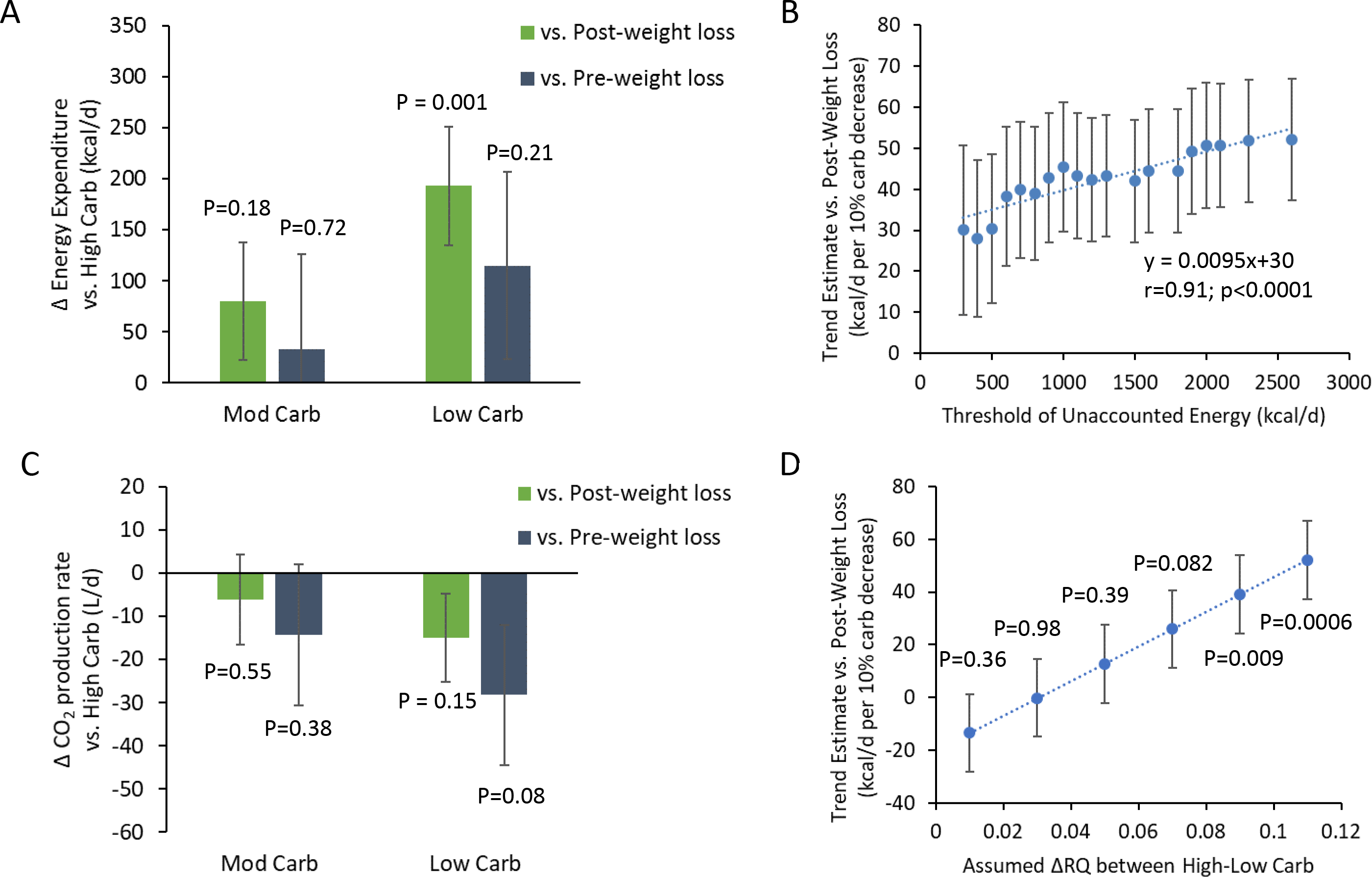
A) Intention to treat analysis of differences in total energy expenditure (TEE) consuming low and moderate carbohydrate diets compared to subjects consuming a high-carbohydrate diet. The green bars illustrate the significant effect of the low carbohydrate diet on average TEE during weight loss maintenance as compared to the immediate post-weight loss period. The gray bars indicate the lack of significant effect of diet on average TEE during weight loss maintenance as compared to the pre-weight loss baseline period. B) Trend estimate for the TEE diet effect during weight loss maintenance (calculated using the revised plan comparing to the post-weight loss TEE) as a function of the threshold used to filter out subjects with excessive relative amounts of unaccounted energy. The rightmost data point includes all 162 subjects with as much as 2600 kcal/d of unaccounted energy and corresponds to the diet effect size reported by Ebbeling et al. according to their revised analysis plan. The leftmost data point indicates a reduced effect size and includes 81 subjects with as much as 300 kcal/d of unaccounted energy. C) Differences in daily average CO_2_ production in the intention to treat group consuming low and moderate carbohydrate diets compared to subjects consuming a high-carbohydrate diet. No significant effects of diet were observed regardless of whether the measurements during weight loss maintenance were compared to the pre-weight loss baseline period (gray bars) or compared to the immediate post-weight loss period (green bars). D) Intention to treat trend estimate for the TEE diet effect during weight loss maintenance (using the revised analysis plan) as a function of the assumed differences in daily RQ between the high and low carbohydrate diets. Error bars are ±SE.

